# Characterizing the middle-age neurophysiology using EEG/MEG

**DOI:** 10.1101/2021.06.04.447084

**Authors:** Justyna Gula, Victoria Moiseeva, Maria Herrojo Ruiz, Marinella Cappelletti

## Abstract

Middle adulthood – the period of life between 40 and 60 years of age – is accompanied by important physical and emotional changes, as well as cognitive and neuronal ones. Nevertheless, middle age is often overlooked in neuroscience under the assumption that this is a time of relative stability, although cognitive decline, as well as changes in brain structure and function are well-established by the age of 60.

Here we characterized the middle-aged brain in the context of healthy younger and older adults by assessing resting-state electrophysiological and neuromagnetic activity in two different samples (N = 179, 631). Alpha and beta oscillations – two key ageing signatures – were analyzed in terms of spectral power and burst events. While posterior alpha power and burst rate features changed linearly with age, similarly to behavioral measures, sensorimotor beta power and burst rate properties varied non-linearly, with inflection points during middle age. The findings suggest that ageing is characterized by distinct spatial and temporal brain dynamics, some critically arising in middle age.

## 1. INTRODUCTION

Middle adulthood – the period of life from about 40 to 60 years old – is often a rich and active stage in most people’s lives, also accompanied by important cognitive and neuronal changes (Lachman, 2004, 2015). Middle age is however often neglected in neuroscience on the account that more informative and larger changes are observable when comparing young to older adults (Lachman, 2010; Willis et al, 2010). Indeed, most studies of the ageing brain are dominated by cross-sectional investigations typically comparing only students (about 20 years old) and pensioners (about 60-80 years old) (Craik & Bialystok, 2006), therefore assuming a linear trajectory of change, with performance in midlife expected to fall midway between young and old age. Non-linear trajectories of age change are rarely tested even when middle agers as part of the sample (Callaghan et al, 2014; Draganski et al, 2011).

In contrast to these studies, cognitive decline is reported to start from mid-age, for instance in terms of impoverished processing speed, memory and executive functions (Ferreira et al, 2015; Gautam et al, 2011; Park et al, 2014; Salthouse, 2009, 2011; Singh-Manoux et al, 2012; Zimprich and Mascherek, 2010). These age-related behavioral changes parallel alterations in brain structure and function, especially in brain regions such as the prefrontal cortex, the medial temporal areas and the hippocampus. These areas show accelerated shrinkage (Raz et al, 2005, 2007, 2010; Fjell et al, 2009), and white matter decreases in volume (Ge et al., 2002) or myelin content starting in the fourth decade of life (Karolis et al, 2019), pointing to the importance of studying middle age for signaling later age-related brain changes.

A complementary approach to previous neuroimaging studies consists of using electroencephalography (EEG) or magnetoencephalography (MEG) to provide a more direct assessment of neurophysiological processes (Cohen, 2017). Neuronal oscillations in EEG/MEG signals can be linked to neurophysiological mechanisms, which is central to understanding age-related brain changes (Sherman et al, 2016, Law et al, 2019). Past M/EEG investigations comparing younger and older adults at rest showed negative correlations between alpha (8-13 Hz) oscillations and age in occipito-parietal brain regions (Ishii et al 2017; Vlahou et al, 2015) or the reverse pattern in beta (13-30 Hz) oscillatory activity in motor areas (Rossiter et al, 2014; Rondina et al, 2019).

Alpha oscillations are known for supporting functional inhibitory mechanisms -including in the context of working memory processing- by suppressing task-irrelevant areas and in turns goal-irrelevant stimuli and behavioral responses (Jensen & Mazaheri, 2010; Klimesch, 2012). Beta oscillations, on the other hand, are prominent in the basal ganglia and in cortical prefrontal and sensorimotor regions (Lundqvist et al, 2016; Schmidt et al, 2019). Although there is an ongoing debate regarding the functions of sensorimotor beta, growing evidence indicates that sensorimotor beta activity is enhanced in association with the maintenance of ongoing motor states, while it decreases during updates in motor plans and predictions (Hosaka et al, 2015; Tan et al, 2016, Sporn et al, 2019).

New insights into the functional role of alpha and beta oscillations can be provided by complementing classical spectral power analysis (collapsing trial information) with the analysis of oscillation “bursts” in single trials (Feingold et al, 2015; Shin et al, 2017; Lundqvist et al, 2016). Brief oscillation bursts of ~50-100 ms duration are a central feature of physiological alpha and beta activity (Poil et al., 2008; Feingold et al, 2015; Torrecillos et al, 2018). In the case of beta bursts, their distribution is maximal in premotor-motor and basal ganglia areas (Feingold et al. 2015; Little et al., 2019). Neurological or psychiatric conditions can modulate spectral power and burst properties (Tinkhauser et al, 2017; Torrecillos et al, 2015; Sporn et al, 2020), yet it remains unclear whether healthy lifespan changes in neuronal oscillations also alter burst properties.

Here we assessed both the spectral power and burst properties of alpha and beta oscillations to capture dynamic changes in neuronal activity that were modulated by ageing. In our first study, EEG was recorded at rest in a group of middle agers as well as in younger and older healthy adults. We hypothesized (i) linear changes with age in alpha activity (Ishii et al 2017; Vlahou et al, 2015), and (ii) non-linear lifespan changes in beta activity, peaking in middle age. To assess the link between alpha oscillations and cognitive performance, we tested whether the hypothesized linear age-related changes in spectral power and bursts properties corresponded with changes in working memory performance, previously shown to modulate alpha activity (Jensen & Mazaheri, 2010; Klimesch, 2012). In a second study, we analyzed a large open MEG dataset for age-related obtained from the Cam-CAN repository (Shafto et al, 2014; Taylor et al, 2017), aiming to replicate the results from our EEG dataset and to determine whether any non-linear age changes in sensorimotor beta activity were paralleled by a similar age-related non-linear modulation of sensorimotor task performance – as available in this dataset.

## 2. METHODS

### 2.1. EEG

#### 2.1.1 Participants

A total of 179 healthy adults provided written consent and received monetary compensation to participate in this study which was approved by the local Ethics Committee at Goldsmiths, University of London. None of the participants had history of neurological or psychiatric disorders, was under regular medication, or showed major cognitive impairments assessed with the Mini-Mental State Examination (MMSE; Folstein et al, 1975; only for participants over 60 years). The sample was divided into 3 age groups: young (mean age 30 ± 5.2; age range 19–39; 61 participants; 41 females), middle age (mean age 48.2 ± 6.2; age range 40–60; 58 participants; 30 females), and older (mean age 69.3 ± 5.5; age range 61–80; 60 participants; 36 females).

#### 2.1.2. Experimental design and procedure

EEG signals were recorded in all 179 healthy participants during 5 minutes of wakeful rest (resting-state EEG, rs-EEG) with eyes open. This was followed by a retro-cue working memory task (see 2.1.4 and Borghini et al, 2018) in 163 of the total participants.

Participants were seated 60 cm from a 21’’ CRT monitor in a darkened room. They were asked to keep their eyes open and focused on a white fixation cross (visual angle 6 degrees) which was programmed and run on ePrime software (Schneider et al, 2002), and displayed on a black screen. Following the rs-EEG recording, participants who took part in the retro-cue proceeded to perform the task (see below). The retro-cue working memory task was programmed in MatLab® v.8.4 (http://www.mathworks.co.uk) using the Cogent toolbox (http://www.vislab.ucl.ac.uk/cogent.php).

#### 2.1.3. EEG acquisition and preprocessing

EEG data were recorded using a 64 channel BioSemi Active Two amplifier system with 64 Ag/AgCl electrodes, sampled at 1024 Hz. The electrooculogram (EOG) was also recorded. Pre-processing was performed using EEGLAB toolbox v2019.0 for MATLAB ® (Delorme & Makeig, 2004) and built-in MATLAB (v. R2019b) function. The EEG signal was re-referenced to the average of two earlobe electrodes and down-sampled to 256 Hz. High-pass (0.5 Hz) and notch (49 - 51 Hz) filters were applied. The data were visually inspected to remove large muscle artifacts. An Independent Component Analysis (ICA; runica) was subsequently used to remove eye-blink and saccadic artifacts. Noisy channels were interpolated using spherical interpolation.

#### 2.1.4. Retro-cue working memory paradigm

We used an established retro-cue working memory (WM) task providing a continuous rather than a binary measure of WM accuracy as well as source of errors (Borghini et al, 2018; Pertzov et al, 2013). Participants memorized a 1000 ms display of four arrow stimuli differing in color and orientation, followed by a 1000 ms delay during which a 100 ms retro-cue (valid, invalid or neutral) may or may not have been presented. A 3000 ms delay period then preceded the presentation of a randomly oriented arrow (the probe) of the same color as one of the arrows in the memory array. Participants used a continuous, analogue response to match it as closely as possible to the remembered orientation with a maximum response time of 3500 ms (see Supplementary Material and <u>Figure S1</u>).

#### 2.1.5. Behavioral Data analysis

The analysis of performance followed the approach described by Borghini et al (2018; see Supplementary Material). The retro-cue task provides a measure of accuracy (recall precision) as well as the source of error, consisting of the probability of reporting a target item (*pT*), the probability of reporting a non-target item (*pNT,* a measure of mis-binding), random answer (*pU*) and Kappa (κ, an index of variability of the answer).

#### 2.1.6. Spectral power analysis

The power spectral density was estimated by Welch’s periodogram with a Hamming window with a length of 1 s and a 50% overlap. Power was normalized into decibels (dB) with the mean power in the 0.5 − 90 Hz band separately for each channel.

#### 2.1.7. Region and frequency of interest: comparison of young and older participants

To define spatial and spectral regions of interest for subsequent analyses, we ran two-sided cluster-based permutation tests between young and older participants on the spectral power in the alpha (8–12 Hz) and beta (13–30 Hz) bands, separately (1000 randomizations: Maris and Oostenveld, 2007). The algorithm implemented in the FieldTrip toolbox (Oostenveld et al, 2011) mitigates issues related to multiple comparisons by controlling the family-wise error rate (FWE). We extracted the electrodes and frequency bins from the identified significant clusters and used them for subsequent analyses of oscillatory activity (channel-average power, channel-average burst distribution slope τ – See 2.1.8) across the lifespan. Statistical analysis of spectral power was performed using FieldTrip v.20191127 and built-in MATLAB functions.

#### 2.1.8. Burst analysis: Matching pursuit

Burst detection was performed using the Matching Pursuit algorithm (MP, Mallat and Zhang, 1993) with Enhanced Matching Pursuit Implementation (empi, https://github.com/develancer/empi, Kuś et al, 2013). MP has high time-frequency resolution and provides detailed parametrization of the signal based on an iterative algorithm that decomposes the signal into linear combination of functions *g*_i_, selected from a pre-defined, redundant set (dictionary *D*) such that ||*g_i_*|| = 1. In each step *n*, the function *g* was selected to maximize the dot product with the residuals obtained in the preceding step, by subtracting function *g_n−1_* from residuals *R^n−1^x.* For details, see Supplementary Methods.

We used the *empi* multichannel algorithm, which has a predefined, optimal dictionary of Gabor functions (Kuś et al, 2013), such that the signal *x* is represented as a set of well-defined Gabor functions. Detailed parametrization of this representation allowed the selection of specific structures of the signal in terms of amplitude, duration, position, frequency, energy, and phase.

We set as criterion for burst detection an amplitude that exceeded the mean amplitude of the signal in each frequency band ± SD. To exclude short spikes and reduce noise, we chose bursts with width equal to at least two cycles of the band oscillation. An example of bursts selection using the MP algorithm is presented in <u>Figure 1A</u>. In the present study, we selected bursts in the high alpha (10-12 Hz) and full beta (13-30 Hz) band because these frequency ranges were identified in the statistical comparison of young and older participants described in 2.1.7. Note that this approach was blind to any effects in middle age, which was of particular interest in our study. The quantitative and statistical analyses of the burst events focused on the duration parameter. This was complemented by a qualitative description of the frequency distribution of bursts within the alpha and beta bands.

**Figure 1.**
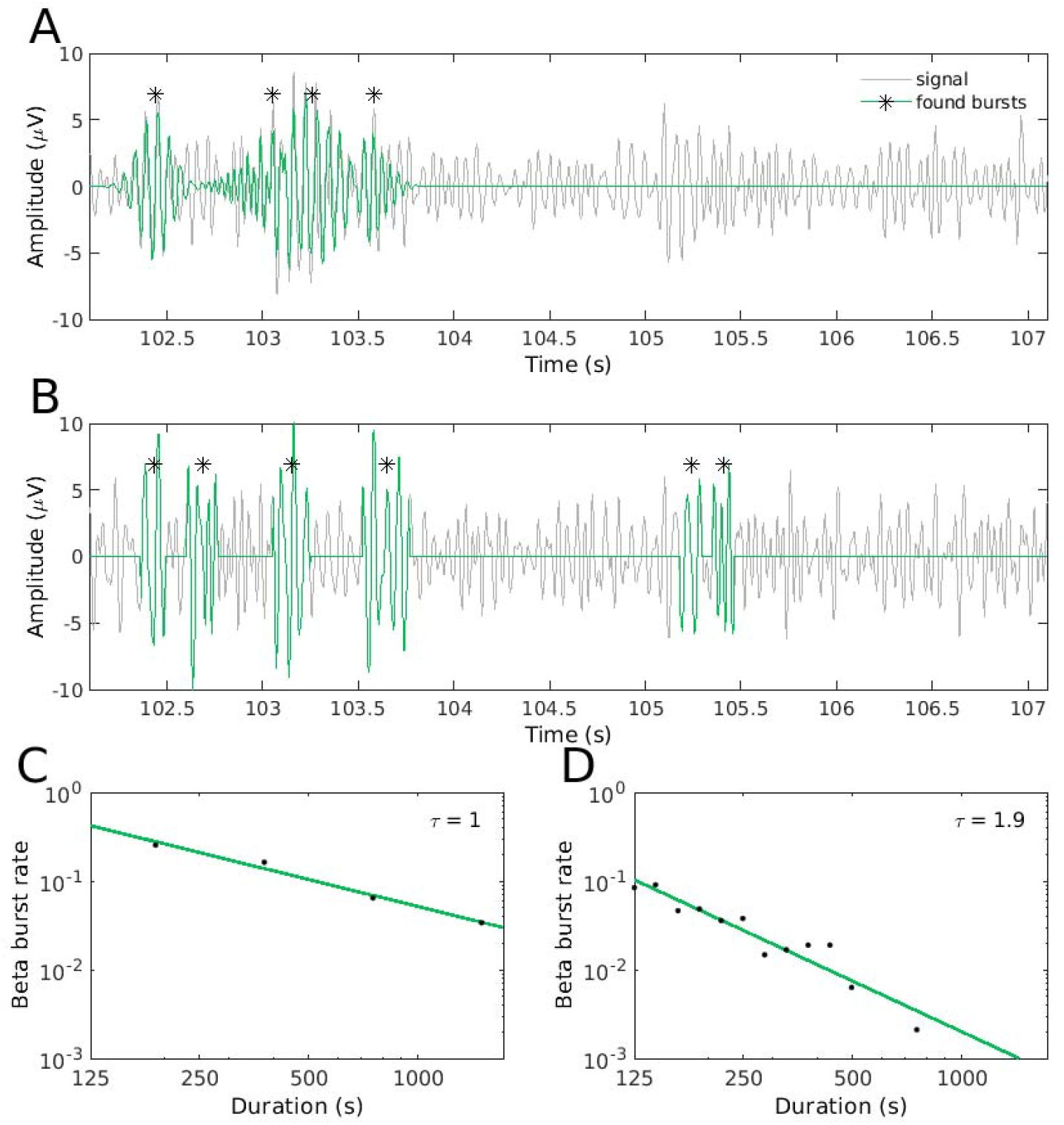
Methods: Matching Pursuit and Amplitude-envelope burst detection. (A) Example of the reconstruction of beta oscillation (grey) and detected bursts (green) in an exemplary participant in C3 channel using the matching pursuit algorithm (MP). Asterisks mark the centre of the burst. (B) Example of beta oscillation (grey) and detected bursts (green) in the same participant and channel as (A) but using the amplitude-envelope method. Asterisks mark the centre of the burst. (C) Distribution of beta bursts per duration in log-log scale in the same participant and channel as (A-B) using the MP method. For each channel, the slope (τ) was extracted. (D) Distribution of beta bursts per duration in log-log scale in the same participant and channel as (A-C) using the amplitude-envelope method. For each channel, the slope, τ, was extracted.

#### 2.1.9. Burst analysis: amplitude envelope

Next, we used a complementary burst detection method based on the oscillations amplitude envelope (Poil et al, 2008; Tinkhauser et al, 2017) to replicate the results obtained with MP. We applied bandpass filters separately at 10-12 Hz and 13-30 Hz in the alpha and beta bands, respectively, in line with the relevant frequency ranges identified by the cluster-based permutation test conducted in young and older participants (See 2.1.7). Following past research (Sporn et al, 2020; Tinkhauser et al, 2017), we calculated the amplitude envelope of the oscillation of interest (beta or alpha range) using the Hilbert transform. We then used the 75% percentile of the amplitude envelope of the given oscillation band as threshold. A signal segment exceeding this threshold was detected as burst, but only if its duration was longer than two cycles of 20 Hz and 10 Hz oscillations for the beta and alpha band, respectively. Amplitude threshold crossings separated by less than two cycles were considered to pertain to the same burst. An exemplary selection of bursts using the envelope method is shown in <u>Figure 1B</u>.

#### 2.1.10. Slope of the burst distribution

After detecting bursts with either method, we computed the slope of the distribution of burst rate against burst duration (Poil et al, 2008; Sporn et al, 2020) and assessed its modulation across the lifespan. In brief, a linear model was fitted to the distribution of burst rate per duration in log-log scales, separately for each channel and oscillation frequency band. The distribution was estimated in 20 equally distanced bins on a logarithmic scale, in a time range up to 2000 ms and with a minimal burst duration of two cycles (See 2.1.8). The slope of the linear model fit is typically negative, denoting a more frequent presence of brief bursts than long bursts, corresponding with a normal physiological state (Poil et al, 2008; Feingold et al, 2015). The absolute value of the slope, termed τ, was used in subsequent analyses. Of note, decreasing values τ reflect a more frequent presence of long bursts, or a long-tailed burst distribution. We averaged τ across regions of interest specific for a given band (cluster effects, see 2.1.7). An outline of the analysis is presented in <u>Figure 1</u>.

<u>Figure 1</u> reveals that burst detection with the above-mentioned methods differs qualitatively. The MP algorithm decomposed complex structures into separate Gabor functions with distinct frequencies. As a result, some bursts may overlap in time. By contrast, in the envelope method these structures may pertain to one burst if the constrains mentioned in Section 2.1.9 are fulfilled. Importantly, however, using both methods the τ slope coefficient correlated significantly across participants within the high alpha band (Spearman’s rank-order correlation, *r_s_*=0.55, *p*=0.00001), but not the beta band (Spearman’s rank-order correlation, *r_s_*=0.06, *p*=0.42). Despite the differences between burst-detection methods, we considered that both methods would reveal a consistent pattern of lifespan changes in τ.

#### 2.1.11. Statistical analysis: changes in neurophysiology across lifespan

Statistical analysis focused on assessing whether neural properties in terms of oscillation power and τ (slope of the burst distribution) change linearly or nonlinearly across the lifespan. In addition, we investigated whether behavioral performance in the retro-cue working memory task may show a similar trend as the neural dynamics across the lifespan. We therefore fit polynomial models up to a third order to EEG power, τ, and working memory performance (averaged over retro-cue conditions); age was used as a predictor and sex as a variable of no interest. We used the likelihood ratio test in order to select the model that best explained the data. In the case a non-linear and quadratic winning model we extracted the local extremum, whereas we obtained the inflection point in the case of a winning cubic model.

### 2.2. MEG

#### 2.2.1. Cam-CAN dataset

Because modulation of sensorimotor beta oscillations has been consistently associated with changes in sensorimotor processing (Kilavic et al, 2013; Tan et al, 2016), we investigated whether any potential non-linear modulation of resting-state beta activity with age may correspond to non-linear changes in motor performance across the lifespan. Accordingly, in a second study we analyzed an open-access dataset acquired at the Cambridge Centre for Ageing and Neuroscience (Cam-CAN; Shafto et al, 2014; Taylor et al, 2017). The Cam-CAN dataset includes MEG data recorded during wakeful rest and also during a simple cued-button pressing task in 631 subjects (age range 18-88, uniformly distributed and gender balanced).

#### 2.2.2. MEG recording and pre-processing

The Cam-CAN MEG data were recorded using 306-channel Vectorview system (Elekta Neuromag, Helsinki), with sampling frequency of 1 kHz and band-pass filter of 0.03−330 Hz. Head position, EOG, and electrocardiogram (ECG) were continuously registered. The MEG recordings lasted 8 minutes 40 seconds, both during eyes-closed wakeful resting state, and during sensorimotor performance. Noise reduction (using temporal signal space separation, tSSS, Taulu et al, 2005), reconstruction of missing or corrupted MEG channels, continuous head motion correction, and a transformation to a common head position were performed by the Cam-CAN group with MaxFilter (Taulu and Simola, 2006).

We used FieldTrip (v 20191127) and MATLAB (R2019b) to apply a high-pass (0.5 Hz) and notch (49-51 Hz) filter to the MEG sensor signals, similarly to the EEG analysis in the first study. Fieldtrip was also used to remove artifacts related to eye blinks, saccades, and the heartbeat, using the FastICA algorithm (*hyperbolic tangent)* on the continuous data. An automated removal of ICs was implemented following Bardouille et al (2019). Subsequent analyses of MEG data focused on magnetometers (102 sensors) exclusively.

#### 2.2.3. Sensorimotor task

In the sensorimotor task (Shafto et al, 2014), participants pressed a button with a right index finger after unimodal or bimodal audio/visual stimuli. The audio stimuli were 300 ms-binaural tones at one of three pre-selected frequencies (300, 600, or 1200 Hz, equal number of trials, pseudo-randomly ordered). The visual stimuli consisted of two checkerboards presented to the left and right of a central fixation for 34 ms. After a practice trial, participants completed 128 trials. In 120 of them, audio and visual stimuli were presented simultaneously (bimodal trial); in the remaining 8, only one stimulus was presented (unimodal trials). Bimodal and unimodal trials were randomly ordered, with an inter-trial interval between 2 and 26 seconds. Here we analyzed response time (RT) changes in the task across the lifespan.

#### 2.2.4. MEG spectral power and bursts detection: amplitude envelope method

Analysis of MEG data followed the same steps as for EEG data. Changes in spectral power across the lifespan were assessed by first averaging the normalized spectral power (in dB) across the sensors pertaining to a cluster of significant differences between older and young participants.

Due to memory requirements, MP could not be used on the large Cam-CAN dataset as it exceeded our technical capability during the Covid-19 pandemic. Instead, the analysis of alpha and beta-band burst distributions in the MEG signal was completed with the amplitude-envelope method (Poil et al, 2008; Tinkhauser et al, 2017). As noted in ****2.1.10****, the two burst extraction methods differ slightly as the MP is an approximation rather than a transformation method, unlike the amplitude-envelope method (Schönwald et al, 2012). In addition, MP may identify separate burst events that are merged into one in the amplitude-envelope method. Despite these differences, however, the polynomial model fitting procedure showed consistent findings in the lifespan modulation of the burst-distribution exponent τ in the EEG and MEG datasets.

## 3. RESULTS

### 3.1. Changes in EEG oscillation properties across lifespan

#### 3.1.1. Linear changes in alpha band properties across lifespan

The analysis of differences between young and older participants in normalized alpha (8–12 Hz) power spectral density revealed one significant, positive cluster in the high alpha band (10–12 Hz, cluster-based permutation test, p=0.022). This effect reflected significantly higher alpha power in young participants compared to older adults, with the most pronounced difference between groups over the posterior region (<u>Figure 2A</u>). For subsequent analyses, we defined as region of interest the electrodes in this significant cluster within the high alpha band. A detailed parametrization of bursts in the MP method allowed us to assess the distribution of bursts per frequency (<u>Figure 2B</u>). This distribution was shifted towards lower frequencies in older participants because of a higher ratio of bursts in 8–9 Hz relative to younger and middle-aged adults. Like in young participants, the middle-agers’ distribution was centered around 10 Hz, although its tail was longer. Middle-aged participants had a noticeable higher ratio of high alpha bursts (11–12 Hz) than younger and older.

**Figure 2.**
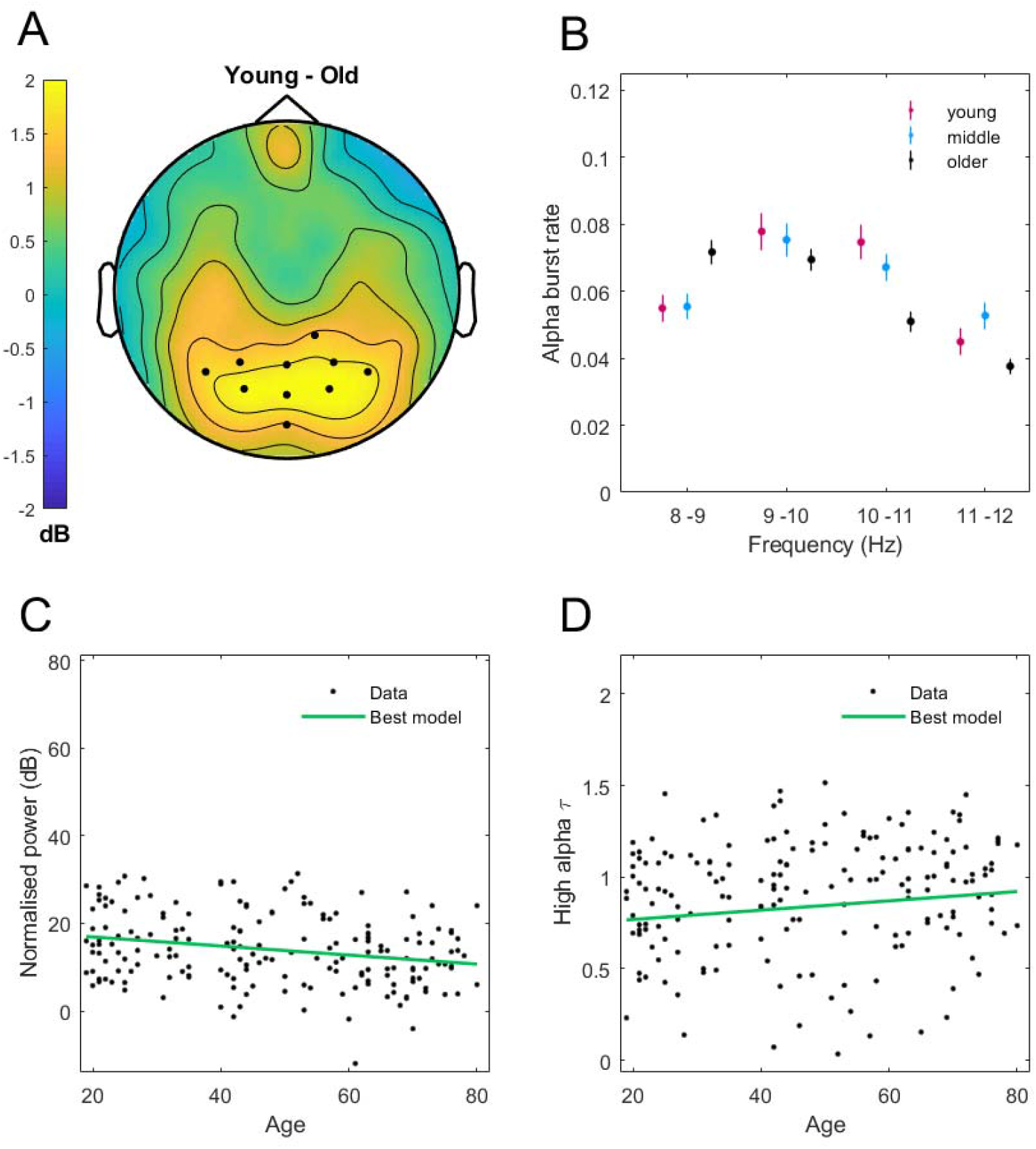
Changes of alpha oscillation across lifespan. (A) In posterior cluster (black dots) young participants showed significantly higher power in the high alpha band relative to older participants. (B) Distribution of bursts per frequency in the three age groups (mean and standard error of the mean). (C) Linear decrease of power in high alpha band across lifespan. (D) Linear increase of τ of high alpha bursts across the lifespan extracted with MP method.

High alpha power in the posterior 10-12 Hz cluster decreased linearly across the lifespan (best model: linear, *p*=0.0054, *R^2^*=0.058), which converged with a linear increase of the exponent τ of high alpha bursts in the same cluster (MP method, best model: linear, *p*=0.016, *R^2^*=0.046, also for the amplitude-envelope method, best model: linear, *p*=1.43·10^−4^, *R^2^*=0.096, see Figure S2). This suggests that with age, the ratio of long bursts within the high alpha band declines as does high alpha power (<u>Figure 2 C–D</u>).

#### 3.1.2. Nonlinear changes in beta band parameters across the lifespan

The cluster-based permutation test showed significant differences in beta power between younger and older adults in the full 13-30 Hz range, and in one negative cluster at central electrodes (*p*=0.001). Younger participants had significantly lower beta power than older adults (<u>Figure 3A</u>). The burst analysis showed that the largest proportion of beta bursts was in the low beta range (13–20 Hz) (<u>Figure 3B</u>). Within this frequency range, qualitative differences between older participants and the other two groups were most visible, with older adults exhibiting more beta bursts (13-17 Hz). In middle-aged adults we observed a higher ratio of bursts within 21–22 Hz than in young and older adults (<u>Figure 3B</u>).

**Figure 3.**
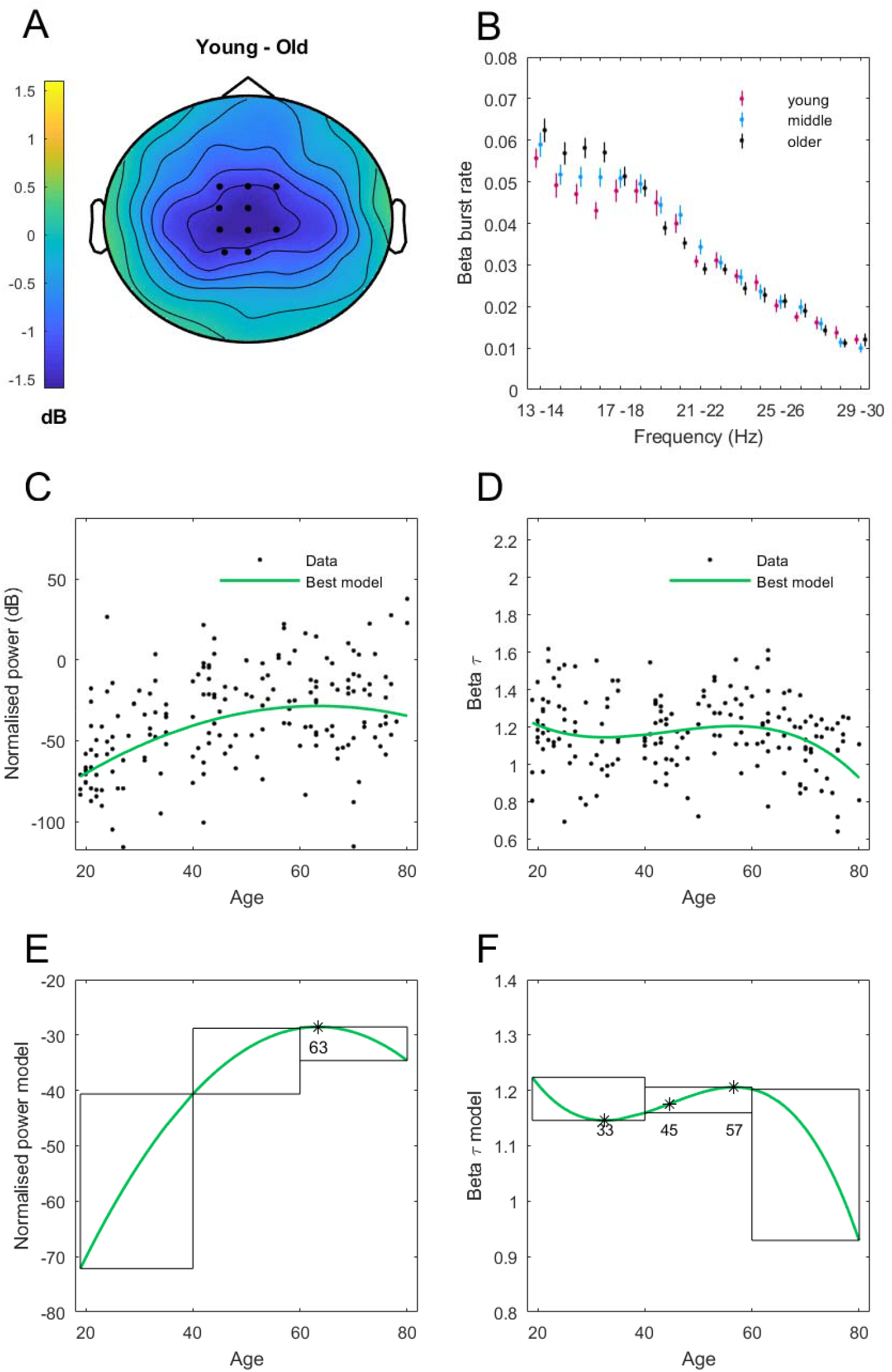
Changes of beta oscillation across the lifespan. (A) In central clusters (black dots) younger participants showed significantly lower power in the beta band compared to older adults. (B) Distribution of bursts per frequency in the three age groups (mean and standard error of the mean). (C) Significant, non-linear changes across lifespan in beta power. (D) Significant, non-linear changes of τ of beta bursts across lifespan. (E) Analysis of the model fitted to changes in beta power. Black lines mark age group ranges. The asterisk indicates the maximum of the function. (F) Analysis of the model fitted to changes in beta τ. Black lines mark age group ranges. Stars indicate maximum, minimum, and inflection point.

The averaged beta power in the significant central cluster varied non-linearly across the lifespan (<u>Figure 3C</u> <u>and</u> <u>E</u>; best model: quadratic, *p*=5.4·10^−10^, *R^2^* =0.23). The initial increase in beta power during youth slowed in middle age and reached its maximum around the age of 60, and slowly decreased in older age. The slope of the burst distribution, τ, extracted with the MP method also changed in a non-linear fashion (<u>Figure 3D</u> <u>and</u> <u>F</u>; best model: cubic, *p*=0.0073, *R^2^* =0.042). At a younger age, τ was high (short-tailed distribution, with a steeper ratio of brief bursts relative to long bursts) and decreased gradually until the age of 30. An increase in τ was then observed in middle age, followed by a steep decline of τ (long-tailed distribution, including more frequent long beta bursts) after the age of 60. These results were not observed using the amplitude-envelope method (all models not significant). Crucially, changes in both beta power and τ (extremum and inflection point) occurred within or close to middle age.

### 3.2. MEG changes across lifespan

#### 3.2.1. Alpha band

In the MEG dataset, cluster-based permutation tests showed significant differences in normalized alpha power between young (N = 160) and older (N = 255) participants (*p* = 0.001). Young participants had higher alpha power than older participants within 10-12 Hz in a region of midline and occipital sensors (<u>Figure 4A-B</u>). Using those sensors and frequency range, we then established that the best model fit to explain the changes in alpha power across the lifespan was linear, demonstrating a significant drop in alpha power with age (<u>Figure 4C</u>; *p* = 4×10^−11^, *R^2^* = 0.11).

**Figure 4.**
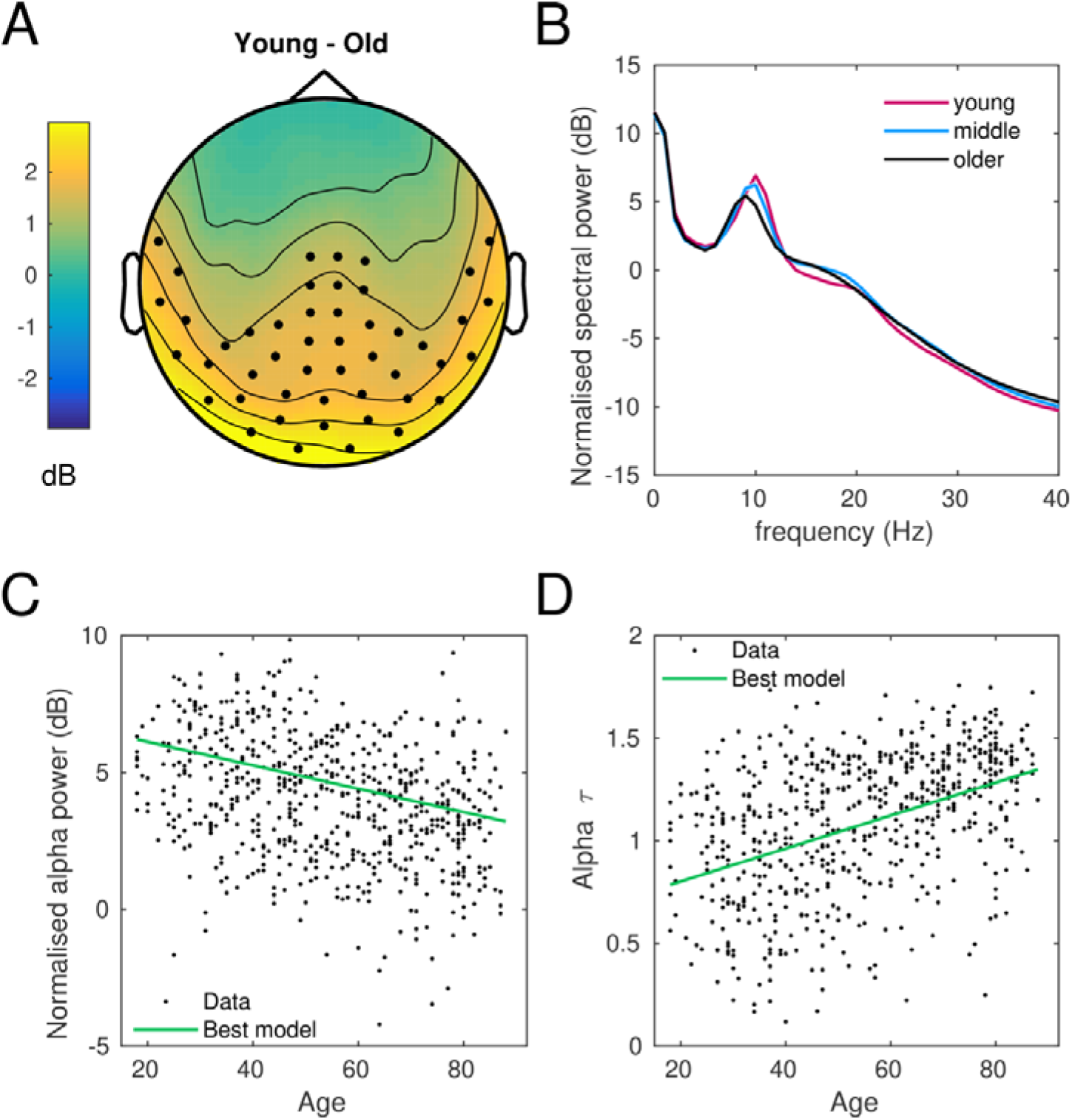
Changes of alpha oscillation across lifespan. (A) In midline-posterior cluster (black dots) younger participants showed significantly higher power in the high alpha band relative to older participants. (B) Normalized spectral power density for young, middle age and older participants. (C) Linear decrease of power in high alpha band across lifespan. (D) Linear increase of τ of high alpha bursts across the lifespan.

In this cluster of sensors and within 10-12 Hz, the slope of the alpha-band burst distribution also changed linearly with age (<u>Figure 4D</u>; *p* = 4×10^−30^, *R^2^* = 0.19). As in the EEG analysis, the slope τ increased linearly with age, suggesting that long-duration bursts were less frequent in older participants, and more frequent in young ones.

#### 3.2.2. Beta band

Converging with the EEG spectral power results, older participants had increased beta power within 13-30 Hz relative to younger ones (p=0.001, cluster-based permutation test). This contrast effect spread centrally (<u>Figure 5A</u>). When averaging beta power within this central cluster, changes in beta power across the lifespan were best described by a non-linear function, also consistent with the EEG results (<u>Figure 5B</u>; quadratic fit, *p*=1 × 10^−7^, *R^2^*=0.10). The maximum of the quadratic fit was found at the age of 58 (<u>Figure 5E</u>, also converging with the EEG results showing a maximum at 63 years old). A cubic function was the best model explaining the modulation of the burst distribution slope, τ, with age (<u>Figure 5C</u>; *p*=0.0057, *R^2^*=0.13). At a younger age, τ decreased steeply, reverting to a slow increase during middle age and followed by an increase-to-decrease transition in older participants (transition at 78 years old; <u>Figure 5E</u>). The lowest slope values of the burst distribution occurred in middle age, suggesting that long-duration beta bursts were more frequent in this age group than in the elderly or young population.

**Figure 5.**
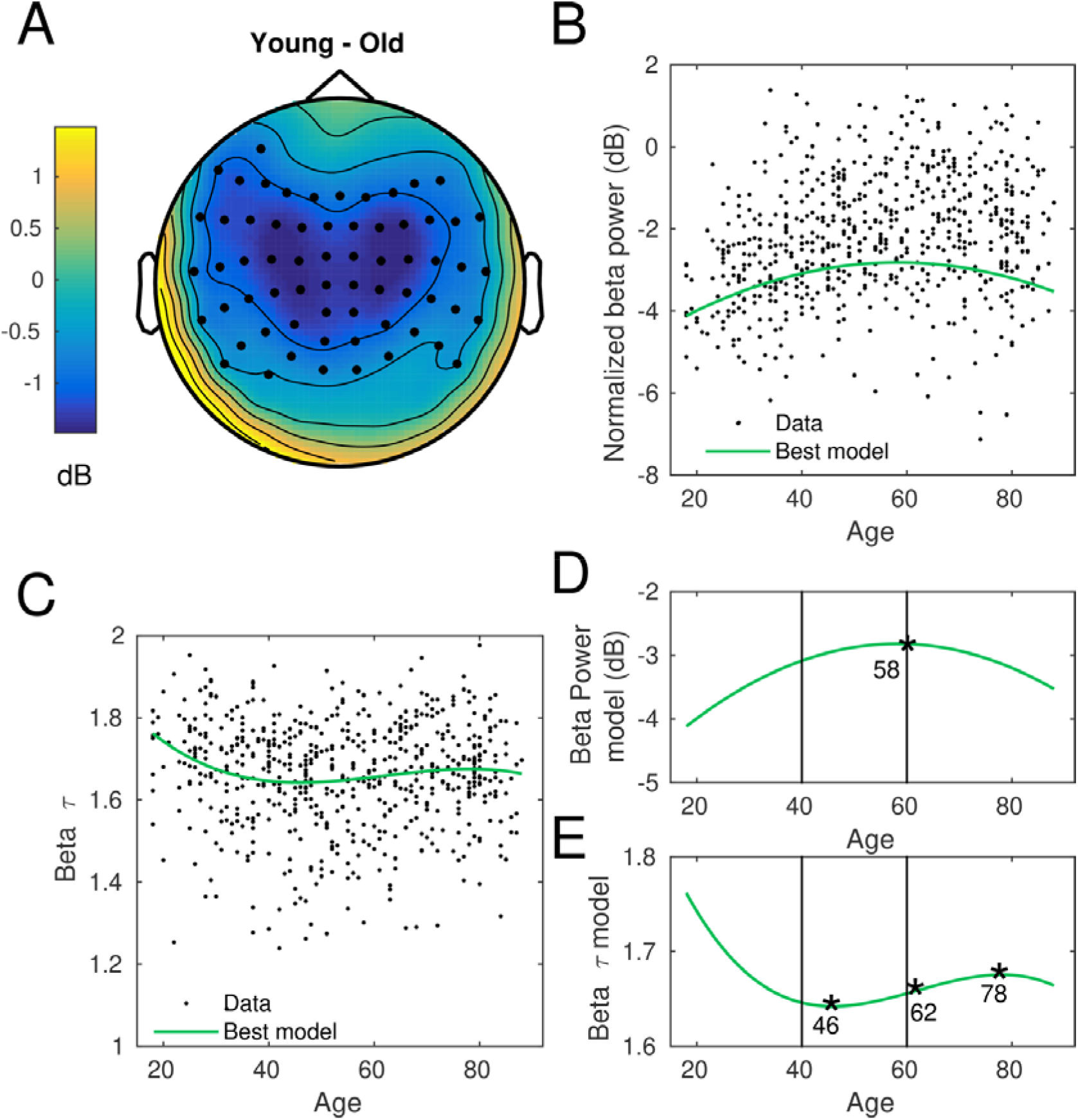
Changes of beta oscillation across lifespan. (A) In central clusters (black dots) younger participants showed significantly lower power in the beta band than older adults. (B) Significant, non-linear changes across lifespan in beta power. (D) Significant, non-linear changes of τ of beta bursts across lifespan. (E) Analysis of the model fitted to changes in beta power. Black lines mark age group ranges. The asterisk indicates the maximum of the function. (F) Analysis of the model fitted to changes in beta τ. Black lines mark age group ranges. Stars indicate maximum, minimum, and inflection point.

### 3.3. Behavioral changes across lifespan

#### 3.3.1. Working Memory Performance

Mean performance across the three retro-cue conditions was analyzed. Recall precision, the probability of reporting the correct item (pT), and variability of the answers (κ) declined linearly as a function of age (precision, *p* = 2.6·10^−5^, *R^2^* = 0.13; pT, *p* = 0.04, *R^2^* = 0.04; κ, *p* = 0.02, *R^2^* = 0.05, <u>Figure S3 A-C</u>; note that a decrease in κ indicates increased variability of the answer). As predicted, the decline in retro-cue working memory performance mirrored the age-related linear changes observed in the alpha band.

#### 3.3.2. Sensorimotor task performance

An analysis of the mean and variance of response times (RT) in the sensorimotor task across the lifespan indicated that both RT mean and variance increased with age. Unlike beta activity measures, however, this increase was linear and significant only for the RT variance index, which assesses the variation of timing performance within the experimental session (*p* = 4×10^−6^, *R^2^* = 0.08, <u>Figure S3 D</u>).

## DISCUSSION

This study investigated the middle-aged brain in the context of younger and older adults by assessing electrophysiological and neuromagnetic responses in alpha and beta oscillations during rest. Neural oscillatory activity was examined in terms of two features – spectral power and burst events. In addition, the possible association between these measures and cognitive and sensorimotor performance was explored. The analysis of spectral power and burst properties based on two complementary methods demonstrated consistent results corresponding to lifespan changes in alpha and beta oscillatory activity in two independent datasets. While alpha power and the slope of the alpha burst distribution changed linearly with age, primarily in posterior brain regions, sensorimotor beta oscillatory power and the burst distribution slope varied non-linearly across the lifespan. These findings suggest that the brain undergoes distinct ageing processes characterized by different spatial and temporal dynamics, some critically arising in middle age.

As predicted, alpha power declined linearly with age and more strongly in occipital clusters. This finding is in line with observations that physiological aging corresponds to gradual changes in spectral power profile, with a pronounced amplitude decrease in alpha power (Ishii et al, 2017; Rossini et al, 2007; Vlahou et al, 2014). Previous results on this linear age-related decrease in alpha power stemmed primarily from binary comparisons of younger and older adults (e.g., Ishii et al 2017; Rossini et al, 2007; Vlahou et al, 2015). Our data extend this evidence, supporting that alpha oscillatory activity declines with age also when middle agers are included in the sample.

Complementing the alpha power results, the analysis of burst events in the alpha band showed that the burst distribution exponents, τ, also changed linearly with age. Thus, young adults had more frequent long-duration alpha bursts than older participants, providing further support for linear changes in alpha activity across the lifespan. Brief alpha bursts are a normal physiological feature of resting state activity (Poil et al, 2008). The distribution of burst duration values typically exhibits a power-law decay, which is a hallmark of scale-free temporal dynamics in human brain oscillations (Poil et al, 2008; Roberts et al, 2015). Compared to beta (and gamma) bursts, alpha bursts have received considerably less attention, and thus our study offers novel insight into the lifespan changes of physiological alpha bursts at rest.

Alpha oscillations are mainly generated by cortico-cortical and thalamo-cortical neuronal networks (Duchanova and Christov 2014). These networks modulate the cholinergic system of the basal forebrain, which is important for regulating high-level cognitive abilities such as attention and memory (Ishii er al, 2017). Linear age-related changes are reported in structures supporting thalamo-cortical connections such as the thalamus (Pfefferbaum et al, 2013), and in alpha-source posterior brain regions (Babiloni et al, 2006; Hindriks & van Putten, 2013; Ishii et al 2017). Cognitive processes relying on these regions, like the retrieval of information, as well as attention are also characterized by linear age-related changes (Steriade and Llinas, 1988; Brunia, 1999). The age-related linear changes we observed in alpha power and τ exponents are therefore consistent with and complement previously reported evidence of linear changes in alpha oscillatory activity (power) in young vs older adults (Babiloni et al, 2006; Haegens et al 2014). Interestingly, the linear changes in alpha-band activity that we observed corresponded to linear changes in working memory performance. An open question for future research is whether alpha burst events may be a more sensitive marker of individual differences in performance of cognitive and working memory tasks than classical occipital alpha power.

A novel result revealed by our data was the non-linear trajectory of beta oscillatory activity across the lifespan, which we obtained independently in the EEG and MEG datasets. Beta power exhibited a quadratic age-related trajectory: an initial increase in beta power in younger adults slowed during middle age, reached its maximum at around the age of 60, and was then followed by a slow decrease in older age. The exponents τ of the beta burst distribution also changed non-linearly with age, albeit in a decreasing fashion and following a cubic function. At a younger age τ was greater, indicating a higher ratio of brief beta bursts. This was followed by a gradual drop in τ, reaching minimum values in middle age at 46, and then slowly increasing again, until 78, when the exponents exhibited a final slow decay trend. Thus, across both our EEG and MEG datasets middle age and, to a lesser extent, older age, were associated with a more frequent presence of long-duration beta bursts, resembling pathological findings in neurological disorders, such as in Parkinson’s disease (Lofredi et al, 2019; Tinkhauser et al, 2017). Critically, our results indicate that middle age is the period of life when the most salient changes in both beta power and burst exponents τ occurred, suggesting that important physiological transformations take place in midlife.

Brief oscillation bursts are considered a healthy feature of beta oscillatory activity in motor cortical areas and in the basal ganglia (Feingold et al, 2015; Little et al, 2019; Poil et al, 2008). A longer duration of beta bursts, however, is an indication of pathological beta rhythms, as shown in the basal ganglia and motor cortex of Parkinson’s disease patients (Lofredi et al, 2019; Tinkhauser, 2017). The occasional presence of longer bursts can also disrupt performance in healthy adults, when occurring before behaviorally relevant cues (Little et al, 2019; Jones, 2016) or during feedback processing (Sporn et al., 2020). Future work should address whether the exacerbated presence of longer bursts at rest in our middle-aged participants is maintained during task performance, and whether it accounts for alterations in behavior and cognition relative to young and old participants.

There is an ongoing debate regarding the functional role of beta oscillations, with some proposals arguing for different types of beta rhythms across different brain regions having different functions (Spitzer and Haegens, 2017; Schmidt et al., 2019). Regarding sensorimotor beta, the classic findings are that it is attenuated during movement and exhibits phasic increases at the end of the action, a phenomenon termed beta rebound and linked to inhibitory GABAergic activity (see Kilavik et al, 2013). More recent work emphasizes its dynamic and parametric nature: flexible moment-to-moment modulations in beta activity contribute to maintaining/updating motor states and predictions through increases/ decreases in power and burst rate, respectively (Hosaka et al, 2015; Tan et al, 2016; Sporn et al, 2020). Thus, increased beta activity and burst rate in cortical-basal ganglia circuits may stabilize motor and also working memory representations –maintaining predictions during ongoing behavior (Bastos et al, 2020; Leventhal et al, 2014; Lundqvist et al, 2016, 2018; Tan et al, 2016; Spitzer and Haegens, 2017).

Of note, in our study, the region of interest for the beta power and burst analyses was obtained from the contrast analysis of old and young groups, which revealed widespread differences at central and anterior electrode regions, a pattern consistent with previous reports (Hari & Salmelin, 1997; Crone et al, 1998). These differences were obtained at rest. Accordingly, the functional significance of the non-linear changes in resting-state beta power and burst exponents with age needs to be specified in future work. This could also shed light on our finding that the independent measure of sensorimotor performance in the MEG dataset changed linearly with age. Along this line, a recent analysis of the CamCan dataset during sensorimotor task performance revealed a linear decrease with age in sensorimotor beta rebound (Bardouille et al, 2019). Exclusively linear models were tested in that study. Our findings suggest that comparing different polynomial models to explain lifespan changes in beta oscillatory activity (power and bursts) during task performance will be a necessary step to clarify whether alterations in beta dynamics are most pronounced in middle-age.

This evidence of specific beta oscillations changes during midlife complements the scarce data on this age group. For instance, past studies showed that relative to younger and older adults, middle age is associated with increased strength (about 10%) of the phase synchronization between beta and alpha oscillations (Nikulin and Brismar, 2006). Middle age also corresponds to reduced power and centre frequency for both slow and fast gamma (Murty et al, 2019), and early work indicated reduced variability of ERP amplitude and of cortical coupling in middle age, which reflect increased homogeneity in the way the brain responds (Dustman et al, 1990; 1993). Middle age is also thought to reflect the optimal balance between inhibition and excitation, set between development and decline (Kosseva et al, 2002; Nikulin and Brismar, 2006; Rossini et al, 2007). Late middle age has been associated to increased homogeneity, which is thought to reflect a shift in the excitatory-inhibitory balance towards reduced inhibition (Dustman et al, 1993). Our results of non-linear trajectory of age-related changes in beta activity complement these observations and reinforce the view that middle age signals important cortical changes relative to younger and older ages.

## ACKNOWLEDGEMENTS

This work was supported in all its aspects by a BIAL Foundation Grant and a British Academy Grant (SG090611). MHR and VM were partially supported by the Basic Research Program at the National Research University Higher School of Economics (Russian Federation).

We thank Chiara Crovace, Mara Golemme, Daniele Porricelli, Elisa Tatti, and Margherita Tecilla for help with data collection.

## Authors’ contributions

MC and MHR: conceptualization, methodology; JG and MHR: data analyses; MC and MHR: supervision of data collection and analyses. MC, MHR and VM: financial support. All authors: writing and approving manuscript.

## REFERENCES

Babiloni C, Binetti G, Cassarino A, Dal Forno G, Del Percio C, Ferreri F, Ferri R, Frisoni G, Galderisi S, Hirata K, Lanuzza B, Miniussi C, Mucci A, Nobili F, Rodriguez G, Romani GL, Rossini P.M. 2006. Sources of cortical rhythms in adults during physiological aging: a multicentric EEG study. HBM. 27:162–172.

Bardouille T, Bailey L, CamCAN Group. 2019. Evidence for age-related changes in sensorimotor neuromagnetic responses during cued button pressing in a large open-access dataset. NeuroImage. 193:25–34.

Bastos A.M, Lundqvist M, Waite A.S, Kopell N, Miller E.K. 2020. Layer and rhythm specificity for predictive routing. PNAS. 117:31459–31469.

Borghini G, Candini M, Filannino C, Romei V, Zokaei N, Walsh V, Hussain M, Cappelletti M. 2018. Alpha oscillations are causally linked to inhibitory abilities in ageing. J Neurosci. 38:4418–4429.

Brunia C.H.M. 1999. Neural aspects of anticipatory behavior. Acta Psychologica, 101, 213–242.

Callaghan M, Freund P, Draganski B, Anderson E, Cappelletti M, Chowdhury R, Diedrichsen J, Fitzgerald T, Smittenaar P, Helms G, Lutti A, Weiskopf N. 2014. Wide-spread age-related changes of the microstructure in the human brain revealed by quantitative MRI. Neurobiol Ageing. 35:1862–72.

Craik F.I, Bialystok E. 2006. Cognition through the lifespan: mechanisms of change. Trends in Cognitive Sciences, 10:131–138.

Crone N.E, Miglioretti DL, Gordon B, Sieracki JM, Wilson MT, Uematsu S, Lesser RP. 1998. Functional mapping of human sensorimotor cortex with electrocorticographic spectral analysis. I. Alpha and beta event-related desynchronization. Brain. 121:2271–99.

Cohen MX. 2017. Where does EEG come from and what does it mean? Trends Neurosci. 40:208–18.

Delorme A, Makeig S. 2004. EEGLAB: an open-source toolbox for analysis of single-trial EEG dynamics including independent component analysis. J Neuroscience Methods. 15:9–21.

Draganski B, Ashburner J, Hutton C, Kherif F, Frackowiak RSJ, Helms G, Weiskopf N. 2011. Regional specificity of MRI contrast parameter changes in normal ageing revealed by voxel-based quantification (VBQ). Neuroimage. 55:1423–34.

Dushanova J, Christov M. 2014. The effect of aging on EEG brain oscillations related to sensory and sensorimotor functions. Advance Medical Science. 59:61–7.

Dustman R.E, Shearer D.E, Emmerson R.Y. 1993. EEG and event-related potentials in normal aging. Progress in Neurobiology. 41:369–401.

Ferreira D, Correia R, Nieto A, Machado A, Molina Y, Barroso J. 2015. Cognitive decline before the age of 50 can be detected with sensitive cognitive measures. Psicothema 27 216–222.

Feingold J, Gibson DJ, DePasquale B, Graybiel AM. 2015. Bursts of beta oscillation differentiate postperformance activity in the striatum and motor cortex of monkeys performing movement tasks. Proc Natl Acad Sci U SA 112:13687–13692.

Folstein M. F. 1975. “Mini-mental state”. A practical method for grading the cognitive state of patients for the clinician. Journal of Psychiatric Research. 12:189–198.

Fjell AM, Walhovd KB, Fennema-Notestine C, McEvoy LK, Hagler DJ, Holland D, Brewer JB, Dale AM. 2009. One-year brain atrophy evident in healthy aging. J Neuroscience. 29:15223–15231.

Gautam P, Cherbuin N, Sachdev PS, Wen W, Anstey KJ. 2011. Relationships between cognitive function and frontal grey matter volumes and thickness in middle aged and early old-aged adults: The PATH Through Life Study. Neuroimage 55: 845–855.

Ge Y, Grossman R.I, Babb J.S, Rabin M.L, Mannon L.J, Kolson D.L. 2002. Age-Related Total Gray Matter and White Matter Changes in Normal Adult BrainPart I: Volumetric MR Imaging Analysis. American J of Neuroradiology. 23:1327–1333.

Hari R, Salmelin R. 1997. Human cortical oscillations: a neuromagnetic view through the skull. Trends Neurosci. 20:44–49.

Haegens S, Händel B.F, Jensen O. 2011. Top–down controlled alpha band activity in somatosensory areas determines behavioral performance in a discrimination task. J Neuroscience. 31:5197–5204

Haegens S, Cousijn H, Wallis G, Harrison P.J. Nobre A.C. 2014. Inter- and intra-individual variability in alpha peak frequency. NeuroImage. 92:46–55.

Hindriks R, van Putten M. 2013. Thalamo-cortical mechanisms underlying changes in amplitude and frequency of human alpha oscillations. NeuroImage. 15:150–63.

Hosaka R, Nakajima T, Aihara K, Yamaguchi Y, Mushiake H. 2015. The suppression of beta oscillations in the primate supplementary motor complex reflects a volatile state during the updating of action sequences. Cer Cortex. 26:3442–52.

Jensen O, Mazaheri A. 2010. Shaping functional architecture by oscillatory Alpha activity: gating by inhibition. Frontiers in Human Neuroscience. 4, doi:.10.3389/fnhum.2010.00186

Jones SR. 2016. When brain rhythms aren’t ‘rhythmic’: implication for their mechanisms and meaning Current Opinion in Neurobiology 40:72–80.

Ishii R, Canuet L, Aoki Y, Hata M, Iwase M, Ikeda S, Nishida K, Ikeda M. 2017. Healthy and Pathological Brain Aging: From the Perspective of Oscillations, Functional Connectivity, and Signal Complexity. Neuropsychobiology. DOI: 10.1159/000486870

Karolis S, Callaghan M, Chieh-En Tseng J, Hope T, Weiskopf N, Rees G, Cappelletti M. 2019. Spatial gradients of healthy aging: a study of myelin-sensitive maps. Neurobiol Ageing.79:83–92.

Kilavik B.E, Zaepffel M, Brovelli A, MacKay W.A, Riehle A. 2013. The ups and downs of beta oscillations in sensorimotor cortex. Experimental Neurology. 245:15–26.

Klimesch W. 2012. Alpha-band oscillations, attention, and controlled access to stored information. TICS. 16:606–17.

Kosseva A.R, Schraderb C, Däuperb J, Denglerb R, Rollnik J.D. 2002. Increased intracortical inhibition in middle-aged humans; a study using paired-pulse transcranial magnetic stimulation. Neurosci Lett 333: 83–86.

Kuś R, Różański PT, Durka PJ. 2013. Multivariate matching pursuit in optimal Gabor dictionaries: theory and software with interface for EEG/MEG via Svarog. BioMed Eng OnLine 12.

Lachman ME. 2004. Development in midlife. Annual Review of Psychology 55:305–331.

Lachman ME, Eileen K, Weiner E.I, Craighead E. 2010. “The Midlife Crisis.” The Corsini Encyclopedia of Psychology 4th Edition, Vol. 3: 993–994.

Lachman ME, Teshale S, Agrigoroaei S. 2015. Midlife as a Pivotal Period in the Life Course: Balancing Growth and Decline at the Crossroads of Youth and Old Age. International J of Behavioral Development. 39:20–31.

Law R.G, Pugliese S, Shin H, Sliva D, Lee S, Neymotin S, Moore C, Jones S.G. 2019. A supragranular nexus for the effects of neocortical beta events on human tactile perception Biorxiv

Leventhal DK, Gage GJ, Schmidt R, Pettibone JR, Case AC, Berke JD. 2012. Basal ganglia beta oscillations accompany cue utilization. Neuron. 73:523–536.

Little S, Bonaiuto J, Bestamann S. 2019. Human motor cortical beta bursts relate to movement planning and response errors. PlosBiology. https://doi.org/10.1371/journal.pbio.3000479.

Lofredi R, Tan H, Neumann WJ, Yeh, C-H, Schneider G.H, Kühn A.A, Brown P. 2019. Beta bursts during continuous movements accompany the velocity decrement in Parkinson’s disease patients. Neurobiology of Disease. 127:462–471.

Lundqvist M, Rose J, Herman P, Brincat SL, Buschman TJ, Miller EK. 2016. Gamma and Beta Bursts Underlie Working Memory. Neuron 90:152–164.

Lundqvist M, Herman P, Warden MR, Brincat SL, Miller EK. 2018. Gamma and beta bursts during working memory readout suggest roles in its volitional control. Nat Commun 9:394.

Mallat S.G, Zhang Z.1993. Matching Pursuits with time-frequency dictionaries. IEEE Trans Signal Process. 41: 3397–3415.

Maris E, Oostenvelt R. 2007. Nonparametric statistical testing of EEG- and MEG-data. J Neurosci Meth. 164:177–90

Murty D, Manikandan K, Kumar W.S, Ramesh R.G, Purokayastha S, Javali M, Prahalada N, Rao N.P. Ray S. 2019. Gamma oscillations weaken with age in healthy elderly in human EEG. NeuroImage. 215, 116826.

Nikulin V.V, Brismar T. 2006. Phase synchronization between alpha and beta oscillations in the human electroencephalogram. Neuroscience. 137:647–657

Oostenveld R, Fries P, Maris E, Schoffelen JM. 2011. Comput Intell Neurosci.. FieldTrip: Open source software for advanced analysis of MEG, EEG, and invasive electrophysiological data. doi: 10.1155/2011/156869.

Park H, Kennedy K.M, Rodrigue K.M, Hebrank, A.C, Park, D.C. 2013. An fMRI study of episodic encoding across the lifespan: changes in subsequent memory effects are evident by middle-age. Neuropsychologia. 51:448–456.

Pertzov Y, Bays P.M, Joseph S, Husain M. 2013. Rapid forgetting prevented by retrospective attention cues. J Experimental Psychology: Human Perception and Performance. 39:1224.

Pfefferbaum A, Rohlfing T, Rosenbloom MJ, Chu W, Colrain IM, Sullivan EV. 2013. Variation in longitudinal trajectories of regional brain volumes of healthy men and women (ages 10 to 85 years) measured with atlas-based parcellation of MRI. NeuroImage. 65:176–193

Poil SS, van Ooyen A, LinkenkaerLHansen K. 2008. Avalanche dynamics of human brain oscillations: relation to critical branching processes and temporal correlations. HBM. 29:770–7.

Raz N, Lindenberger U, Rodrigue KM, Kennedy KM, Head D, Williamson A, Dahle C, Gerstorf D, Acker JD. 2005. Regional brain changes in aging healthy adults: general trends, individual differences, and modifiers. Cer Cortex.15:1676–1689.

Raz N, Rodrigue KM, Kennedy KM, Acker JD. 2007. Vascular health and longitudinal changes in brain and cognition in middle-aged and older adults. Neuropsychology. 21:149–157.

Raz N, Ghisletta P, Rodrigue KM, Kennedy KM, Lindenberger U. 2010. Trajectories of brain aging in middle-aged and older adults: Regional and individual differences. Neuroimage. 51:501–511.

Roberts JA, Boonstra TW, Breakspear M. 2015. The heavy tail of the human brain. Curr Opin Neurobiol. 1;31:164–72.

Rondina Ii R, Olsen R.K, Li L, Meltzer J.A, Ryan J.D. 2019. Age-related changes to oscillatory dynamics during maintenance and retrieval in a relational memory task. PloSOne. 14:e0211851.

Rossini PM, Rossi S, Babiloni C, Polich J. 2007. Clinica neurophysiology of aging brain: from normal aging to neurodegeneration. Progress in Neurobiology. 83:375–400.

Rossiter HE, Davis EM, Clark EV, Boudrias MH, Ward NS. 2014. Beta oscillations reflect changes in motor cortex inhibition in healthy ageing. Neuroimage. 91C:360–365.

Salthouse T.A. 2009. When does age-related cognitive decline begin? Neurob Aging. 30:507–514.

Salthouse T.A. 2011. Neuroanatomical substrates of age-related cognitive decline. Psychol Bulletin. 137:753–784.

Schmidt R. Herrojo Ruiz M, Kilavik B.E. Kilavik BE, Lundqvist M, Starr P. 2019. Beta oscillations in working memory, executive control of movement and thought, and sensorimotor function. J Neurosci. 39:8231–8238.

Schneider W, Eschman A, Zuccolotto A. 2012. E-Prime User’s Guide. Pittsburgh: Psychol Software Tools, Inc.

Schönwald J. S, Carvalho D, de Santa-Helena E, Lemke N, L Gerhardt G, 2012. Topography-specific spindle frequency changes in obstructive sleep apnea. BMC Neurosci, 13: 89.

Shafto et al., 2014; Cam-CAN repository (available at http://www.mrc-cbu.cam.ac.uk/datasets/camcan/

Sherman MA, Lee S, Law R, Haegens S, Thorn CA, Hamalainen MS, Moore CI, Jones SR. 2016. Neural mechanisms of transient neocortical beta rhythms: Converging evidence from humans, computational modeling, monkeys, and mice. Proc Natl Acad Sci U SA 113:E4885–4894.

Shin H, Law R, Tsutsui S, Moore CI, Jones SR. 2017. The rate of transient beta frequency events predicts behavior across tasks and species. Elife 6.

Singh-Manoux A, Kivimaki M, Glymour M.M, Elbaz A, Berr C, Ebmeier K.P, Ferrie J.E, Dugravotet A. 2012. Timing of onset of cognitive decline: results from Whitehall II prospective cohort study. BMJ. 344:d7622.

Spitzer B, Haegens S. 2017. Beyond the status quo: a role for beta oscillations in endogenous content (re) activation. Eneuro. 4(4).

Sporn S, Hein T, Herrojo Ruiz M. 2018. Bursts and variability of beta oscillations mediate the effect of anxiety on motor exploration and motor learning. bioRxiv, :442772.

Sporn S, Hein T, Herrojo Ruiz M. 2020. Alterations in the amplitude and burst rate of beta oscillations impair reward-dependent motor learning in anxiety. eLife. DOI: 10.7554/elife.50654.sa2

Steriade M, Llinas R.R. 1988. The functional states of the thalamus and the associated neuronal interplay. Physiological Review. 68:649–742

Taulu S, Simola J, Kajola M. 2005. Applications of the signal space separation method. IEEE Trans. SignalProcess.533359.

Taulu S, Simola J. 2006. Spatiotemporal signal space separation method for rejecting nearby interference in MEG measurements. Phys Med Biology. 51:1759–68

Tan H, Wade C, Brown P. 2016. Post-movement beta activity in sensorimotor cortex indexes confidence in the estimations from internal models. Journal of Neuroscience. 3:1516–28.

Taylor et al., 2017 Cam-CAN repository (available at http://www.mrc-cbu.cam.ac.uk/datasets/camcan/

Tinkhauser G, Pogosyan A, Tan H, Herz DM, Kuhn AA, Brown P. 2017. Beta burst dynamics in Parkinson’s disease OFF and ON dopaminergic medication. Brain 140:2968–2981.

Torrecillos F, Tinkhauser G, Fischer P, Green A.L, Aziz T.Z, Foltynie T, Limousin P, Zrinzo L, Ashkan K, Brown P, Tan H. 2018. Modulation of Beta Bursts in the Subthalamic Nucleus Predicts Motor Performance. J Neurosci. 38: 8905–8917.

Torrecillos F, Alayrangues J, Kilavik BE, Malfait N. 2015. Distinct Modulations in Sensorimotor Post-movement and Fore-period beta-Band Activities Related to Error Salience Processing and Sensorimotor Adaptation. J Neurosci 35:12753–12765.

Vlahou E.L, Thurm F, Kolassa I.T, Schlee W. 2014. Resting-state slow wave power, healthy aging and cognitive performance. Scientific Report. 4:5101. 10.1038/srep05101.

Willis S.L, Martin M, Rocke C. 2010. Longitudinal perspectives on midlife development: stability and change. European Journal of Ageing. 7:131–134.

Zimprich D, Mascherek A. 2010. Five views of a secret: Does cognition change during middle adulthood? European Journal of Ageing. 7:135–146.

Yu Y, Sanabria D.E, Wang J, Hendrix C.M, Zhang J, Nebeck S.D., Amundson A.M, Busby Z.B, Bauer D.L, Johnson M.D, Johnson L.A, Vitek J.L. 2021. Parkinsonism Alters Beta Burst Dynamics across the Basal Ganglia–Motor Cortical Network. J Neuroscience. 41:2274–2286

