## Supplementary Material for "Characterizing the middle-age neurophysiology using EEG/MEG"

1. **SUPPLEMENTARY METHODS**

***1.2. Retro-cue working memory task***

Our WM paradigm is more complex than most commonly used WM tasks, such as the digit or letter span, because: (1) it allows a more accurate measure of the fidelity or quality of WM representations since it provides a continuous rather than a binary measure of WM performance (Alvarez and Cavanagh, 2004; Bays and Husain, 2008; Bays et al., 2009; Ma et al., 2014); (2) it measures WM accuracy as well as the source of errors; and (3) some of these errors (specifically the probability of target responses and misbinding), reflect inhibitory processes which are linked to alpha oscillations, a key feature of our investigation.

The arrow stimuli (visual angle: 2° x 0.3°) were simultaneously presented to the left and right (2 for each side) of a black, 0.8° diameter fixation cross. Within a trial, they appeared in four of five randomly selected and easily distinguishable colors (yellow, red, blue, green, and white), and were arbitrarily oriented with a minimum of 10° difference between each other. Participants were asked to keep in mind both the orientation and color of these arrows. Each memory array was followed by a 1000 ms delay during which a retro-cue may or may not have been presented (100 ms), by a 3000 ms delay period and finally by the presentation of a randomly oriented arrow (the probe) of the same color as one of the arrows in the memory array. Participants used a continuous, analogue response to match it as closely as possible to the remembered orientation with a maximum response time of 3500 ms (Pertzov et al, 2013; Figure S1).

A total of 126 trials was used, 30% of which (N=38) comprised a neutral cue presented during the memory delay; neutral cues consisted of a white fixation cross that did not change in color. In the other 70% of the trials (N=88), the stimulus display was followed by a 1000 ms delay and by the presentation of an informative retro-cue (for 100 ms). This informative retro-cue indicated the color of the stimulus arrow most likely to be later probed. Within the 70% trials with an informative retro-cue, 70% (N=62) corresponded to the item that was subsequently probed (valid condition). The remaining 30% of the informative retro-cue trials (N=26) consisted of items that were subsequently invalidly probed (invalid condition).

***1.2 Behavioural data analysis***

Accuracy –or recall precision– was defined as the inverse of the circular standard deviation (SD) of the response error, calculated as the angular difference between a participant’s orientation of each arrow stimulus and its veridical orientation in the initial display.

The sources of the errors were estimated following the probabilistic model:

1. $p(\hat{\Theta})=\alpha\phi_{k}(\hat{\Theta}-\Theta)+\beta\frac{1}{m}(\hat{\Theta}-\varphi_{i})+\gamma\frac{1}{2\pi}$

where $\hat{\Theta}$ is the subject’s answer in the trial, Θ is the correct orientation of the target, $\phi_{k}$ is the von Mises distribution with mean equal to 0 and concentration Kappa, $\kappa$. Higher $\kappa$ reflects lower variability of recall target. α and β stand for the probability of responding to the correct item (*pT*) and the probability of responding to a non-target item (*pNT*) respectively. The probability of random answer (*pU*) was defined as $\gamma=1-\alpha-\beta$.

Outliers were identified as scores exceeding mean ± 3SD of recall precision; lower than mean - 3SD for pT; higher than mean + 3SD for pNT, pU, and κ in each condition separately and removed.

***1.3. Burst analysis: Matching pursuit***

Matching Pursuit has been successfully used for detecting sleep spindles (Żygierewicz et al, 1999; Schönwald et al, 2012), event-related analysis of movement-related desynchronization/synchronization (ERD/S; Durka et al, 2001), and tracking seizures (Franaszczuk et al, 1998; Koubeissi et al, 2009).

The algorithm is defined as:

2.

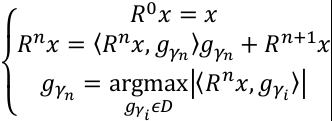

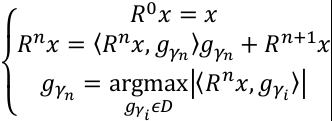

where *R^n^x* is the residuals in *n* step and *x* is the signal. In each step *n*, the function *g* was selected to maximize the dot product with the residuals obtained in the preceding step, by subtracting function
*g_n-1_* from residuals *R^n-1^x.*

We used the empi multichannel algorithm, which has a predefined, optimal dictionary of Gabor functions (Kuś, Różański, Durka 2013):

3.

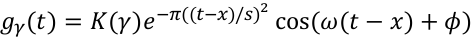

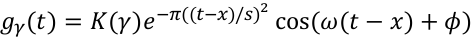

where and are such that In each step, the selected multichannel version of the algorithm finds functions that differ in amplitude only across channels.

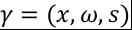

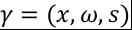

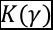

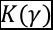

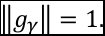

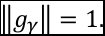

Duration of burst *s* in equation 3 was defined as where σ is the standard deviation of the Gaussian distribution. To ignore short spikes and reduce noise, we chose bursts with width equal to at least two cycles of the band oscillation:, where *freq* was equal to 10 and 20 Hz for the alpha and beta band, respectively.

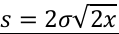

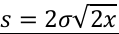

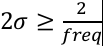

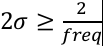

**2. SUPPLEMENTARY FIGURES**

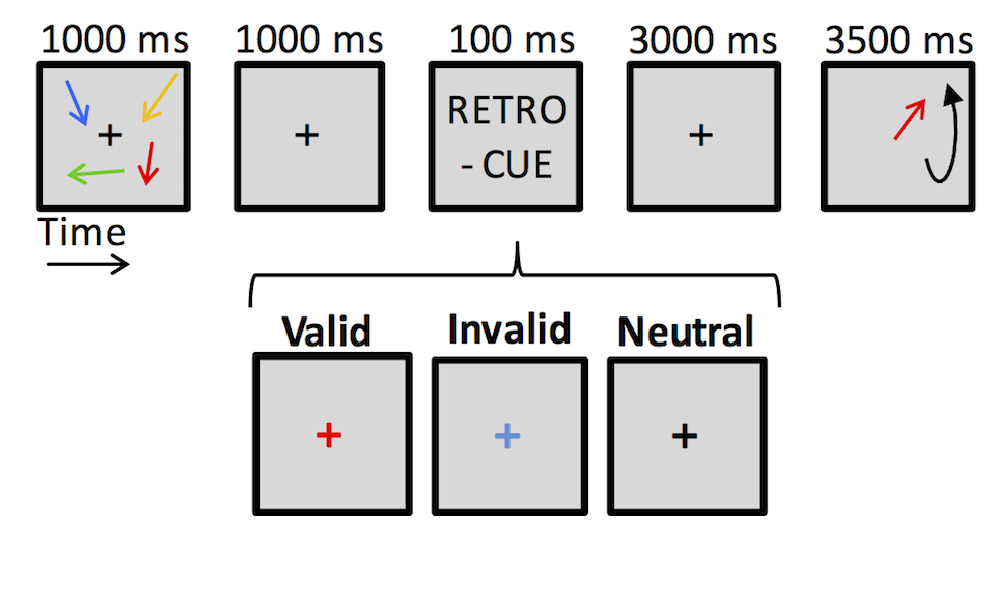

**Figure S1.** The retro-cue working memory task.

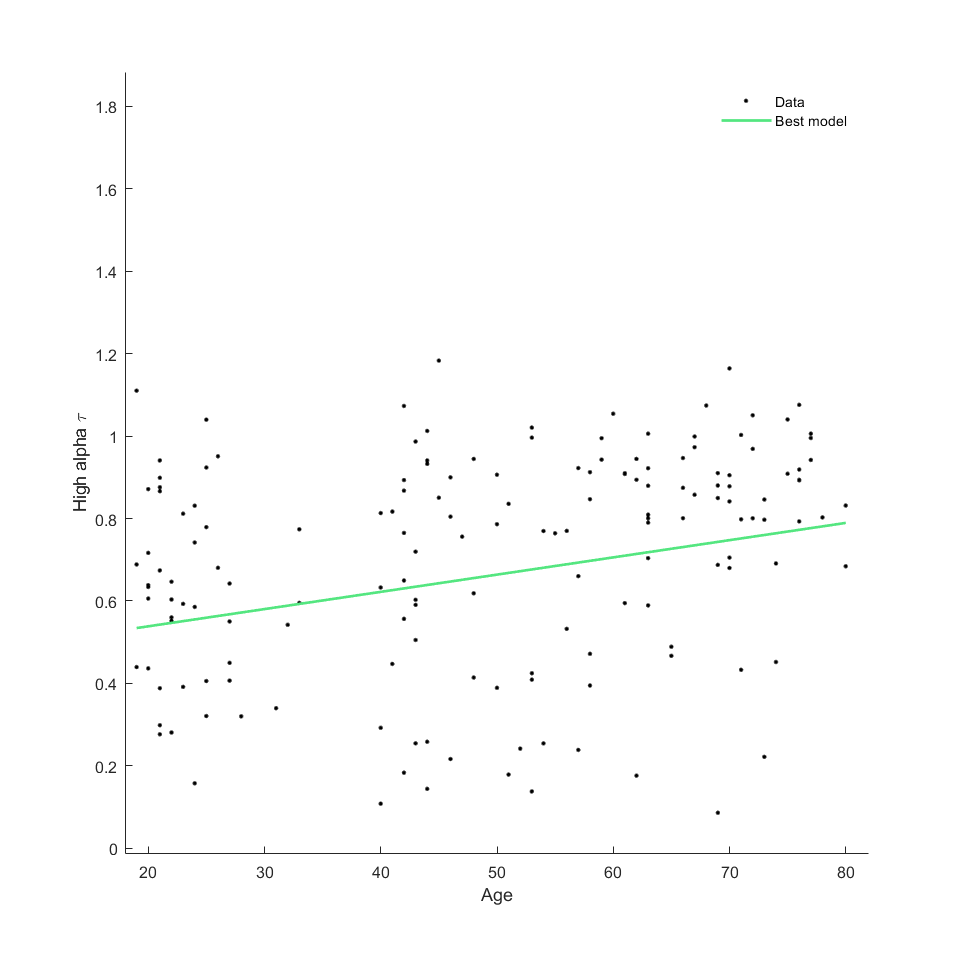

**Figure S2.** Linear increase of τ of high alpha bursts in EEG across the lifespan extracted with amplitude-envelope method.

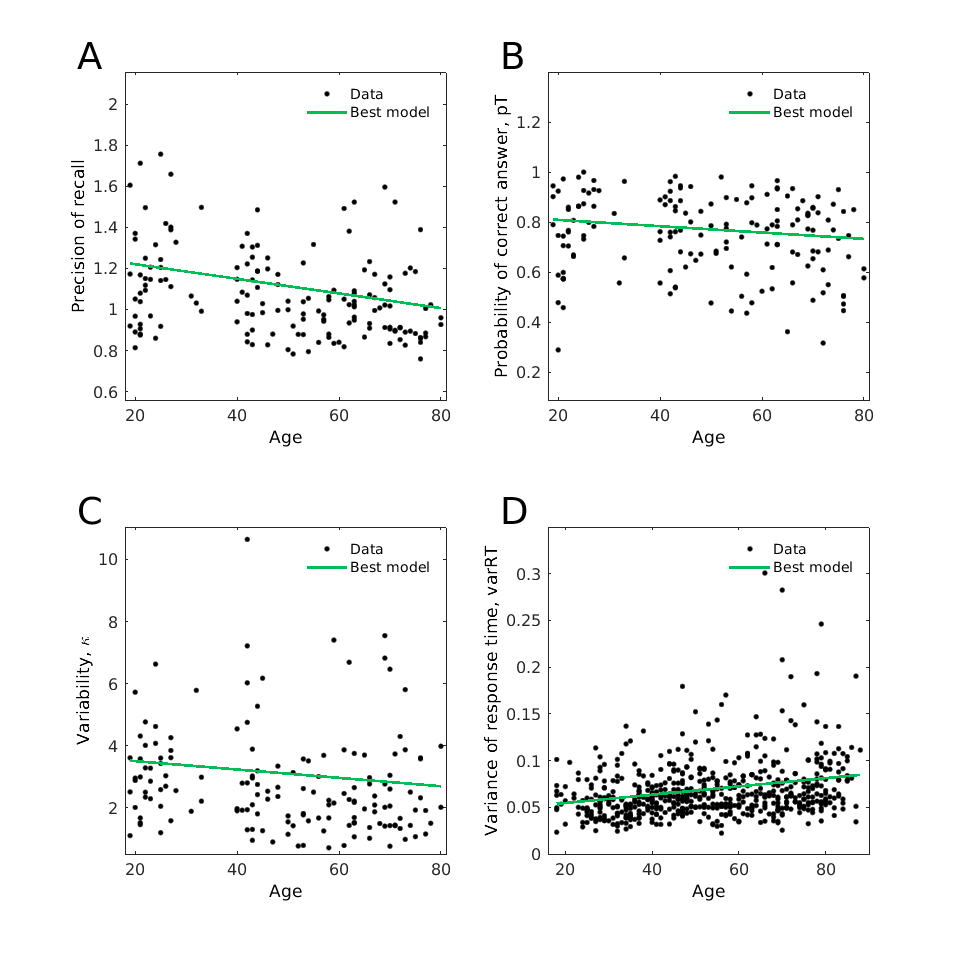

**Figure S3. Changes of retro-cue working memory and sensorimotor tasks performance across the lifespan.** (A) Linear decrease in precision. (B) Linear decrease in probability of reporting target item, pT. (C) Linear decrease in κ (increase of variability of recall target). (D) Linear increase in variance of response time in the sensorimotor task.

**SUPPLEMENTARY REFERENCES**

Alvarez GA. Cavanagh P. 2004. The capacity of visual short-term memory is set both by visual information load and by number of objects. Psychological Science. 15:106-11.

Bays PM, Husain M. 2008. Dynamic shifts of limited working memory resources in human vision. Science. 321:851–854.

Bays PM, Catalao RFG, Husain M. 2009. The precision of visual working memory is set by allocation of a shared resource. Journal of Vision. 9:1-11.

Durka P.J., Ircha D., Neuper C., Pfurtscheller G. 2001. Time-frequency microstructure of event-related electro-encephalogram desynchronisation and synchronisation. Med. Biol. Eng. Comput. 39, 315–321

Franaszczuk, P.J., Bergey, G.K., Durka, P.J., Eisenberg, H.M., 1998. Time–frequency analysis using the Matching Pursuit algorithm applied to seizures originating from the mesial temporal lobe. EEG Clin Neurophysiology, 106: 513-521.

Koubeissi MZ, Jouny CC, Blakeley JO, Bergey GK. 2009. Analysis of dynamics and propagation of parietal cingulate seizures with secondary mesial temporal involvement. Epilepsy Behav. 14:108–112.

Żygierewicz J, Blinowska KJ, Durka PJ, Szelenberger PJ, Niemcewicz S, Androsiuk W. 1999. High resolution study of sleep spindles Clin Neurophysiology. 110:2136-2147.
